# Deep learning for cancer type classification

**DOI:** 10.1101/612762

**Authors:** Zexian Zeng, Chengsheng Mao, Andy Vo, Janna Ore Nugent, Seema A Khan, Susan E Clare, Yuan Luo

## Abstract

Genetic information is becoming more readily available and is increasingly being used to predict patient cancer types as well as their subtypes. Most classification methods thus far utilize somatic mutations as independent features for classification and are limited by study power. To address these limitations, we propose DeepCues, a deep learning model that utilizes convolutional neural networks to derive features from DNA sequencing data for disease classification and relevant gene discovery. Using whole-exome sequencing, germline variants and somatic mutations, including insertions and deletions, are interactively amalgamated as features. In this study, we applied DeepCues to a dataset from TCGA to classify seven different types of major cancers and obtained an overall accuracy of 77.6%. We compared DeepCues to conventional methods and demonstrated a significant overall improvement (p=8.8E-25). Using DeepCues, we found that the top 20 genes associated with breast cancer have a 40% overlap with the top 20 breast cancer genes in the COSMIC database. These data support DeepCues as a novel method to improve the representational resolution of both germline variants and somatic mutations interactively and their power in predicting cancer types, as well the genes involved in each cancer.

## INTRODUCTION

Cancer is a highly complex genetic disease (1,2). Significant mutations termed as driver mutations confer a proliferative advantage in cell growth (2). Advancement in DNA sequencing technologies have led to the systematic processing and analysis of genomes from tumors varying in both cancer type and subtype (3–5). These datasets have resulted in the discovery of relevant functional mutations that affect genes and pathways important in multiple cancers (6–8).

There are patients with unknown primary cancer, where the site of origin cannot be established in the examination of metastatic cancer cells (9). Accurate pathogenetically distinct tumor type classification and accurate site of origin prediction can help identify therapy target precisely and minimizes toxicity and maximizes treatment efficacy (10). In most cases, cancer classification has relied on tumor morphology and histology and gene expression profiling (10–13). Classification based on morphological appearance can be challenging given that tumors with similar histological appearance have been found to respond differently to the same treatment (14). Developing systematic and unbiased approaches for recognizing tumor type and subtype remains essential.

More recently, cancer classification methods have utilized mutational profiles. Somatic mutations were employed to distinguish 28 cancer types resulting in an overall accuracy of 49.4% suggesting somatic point mutations alone as individual variables may not be sufficient to classify cancer types (15). In particular, insertions and deletions as well as germline variation have also been found to be associated with cancer (16,17). However, it has been difficult to integrate germline variation and non-point mutations for cancer-type classification – especially in an interactive manner. Due to the limitation of analysis power, methods including Bayesian classifier (18), regression models (19), and KNN (20) are not optimal in handling high-dimensional features and studying features interactively. In order to circumvent these challenges, labor intensive feature engineering must be done prior to classification that still often lead to poor results (21). Conventional learning algorithms rely heavily on data representations and are typically designed by domain experts. The complexity of the human genome and the amount of required human effort makes it difficult to scale cancer classification studies properly, whereas deep learning can automatically learn a good feature representation (22). Deep learning has recently emerged with the advances in big data, the power of parallel computing and sophisticated algorithms. Furthermore, deep learning models are exponentially more efficient than conventional models in learning intricate patterns from high-dimensional raw data with little guidance (22–25). Typically, convolutional neural networks (CNNs) computes convolution on small regions by sharing parameters between regions (26), which allows training models on large DNA sequences.

Recent work has explored the application of CNNs using raw DNA sequence without defining customized features. DeepBind (27) was proposed to predict specificities of DNA and RNA binding proteins using raw sequence reads, and was reported to outperform the state-of-the-art methods at the time. DeepSEA (28) is a CNN-based tool to learn a regulatory sequence code from chromatin-profiling data, and to predict noncoding variants’ functional effects. DanQ (29) is a hybrid convolutional and bi-directional long short-term memory recurrent neural network for non-coding function prediction; the study reported that there is a 50% precision-recall relative improvement compared with the related models in the area. DeepCpG (30) is another tool based on CNNs to predict methylation states from low-coverage single-cell methylation data; the study reported that methylation states and sequence motifs associated with changes in methylation levels were accurately identified.

Inspired by the successful applications of deep learning models in genomics data, we propose a model to utilize deep learning for disease classification using exome sequencings (DeepCues). Specifically, a CNNs model to study DNA sequences for cancer type prediction is investigated. In addition to tumor sequence, we also investigated whether germline DNA sequence is informative for cancer classification. Furthermore, we are also interested in identifying a subset of genes that are most relevant for each cancer type. In a pilot study utilizing 4,174 samples across seven major cancer types from the Cancer Genome Atlas (TCGA), we were able to achieve an accuracy of 77.6% in predicting seven cancer types. As germline variants dominant somatic mutations number-wise, using only germline information from the same genes, we were able to achieve an accuracy of 73.9%.

## MATERIALS AND METHODS

### Data sources and data processing

Germline and somatic mutations from 4,174 samples across seven major cancer types were obtained from the TCGA (3). The following cancers were analyzed: brain cancer, breast cancer, colorectal cancer, kidney cancer, lung cancer, prostate cancer, and uterus cancer. To obtain germline variants, aligned sequencing data from blood or adjacent normal tissues were recalibrated, and variants were called using HaplotypeCaller in GATK package using assembly hg19 (33). SnpEFF was used for functional annotation (34). Variants annotated with moderate effects were defined as missense mutations and in-frame shifts and variants annotated as high effects were defined as nonsense mutations. Somatic mutations from matched samples were obtained directly from TCGA. In total, 4,600 virtual machines were utilized for 119,000 CPU hours to complete these tasks. In total, we identified 45,119,052 germline mutations and 957,115 somatic mutations from the 4,174 matched samples (Table 1).

**Table 1:**
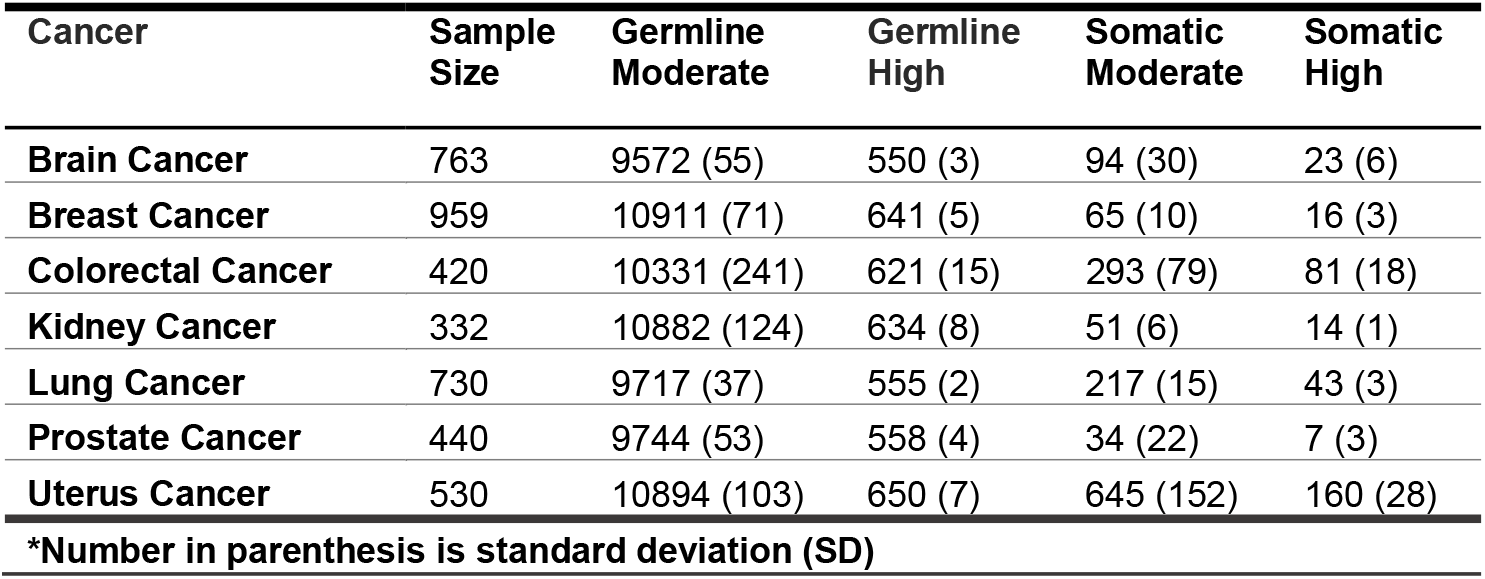
The number of samples of each cancer and the corresponding number of germline variants and somatic mutations. Variants annotated with moderate effects are defined as missense mutations and in-frame shifts; variants annotated as high effects are defined as nonsense mutations.

### Feature generation

Recently, the gene coding direction has been described to be informative for mutational profiles (31). Sequences were annotated as positive if the gene is encoded on the reference strand and as negative if on the complementary strand. To account for directionality, gene sequences used for feature generation were obtained from the RefSeq database (32). Sequences in RefSeq database was started as references, with the ones without mutations in our cohort excluded (Figure 1A). We call these combined reference sequences as consensus matrix. This consensus matrix consists of 24,286 transcripts. The average length of these sequences were 3,375 bases. For individuals, the germline variants identified in the matched normal were constructed into the consensus matrix, forming a germline matrix (Figure 1B). Once a germline matrix was formed for individuals, somatic mutations were then constructed in the germline matrix, forming a germline and somatic matrix (Figure 1C).

**Figure 1:**
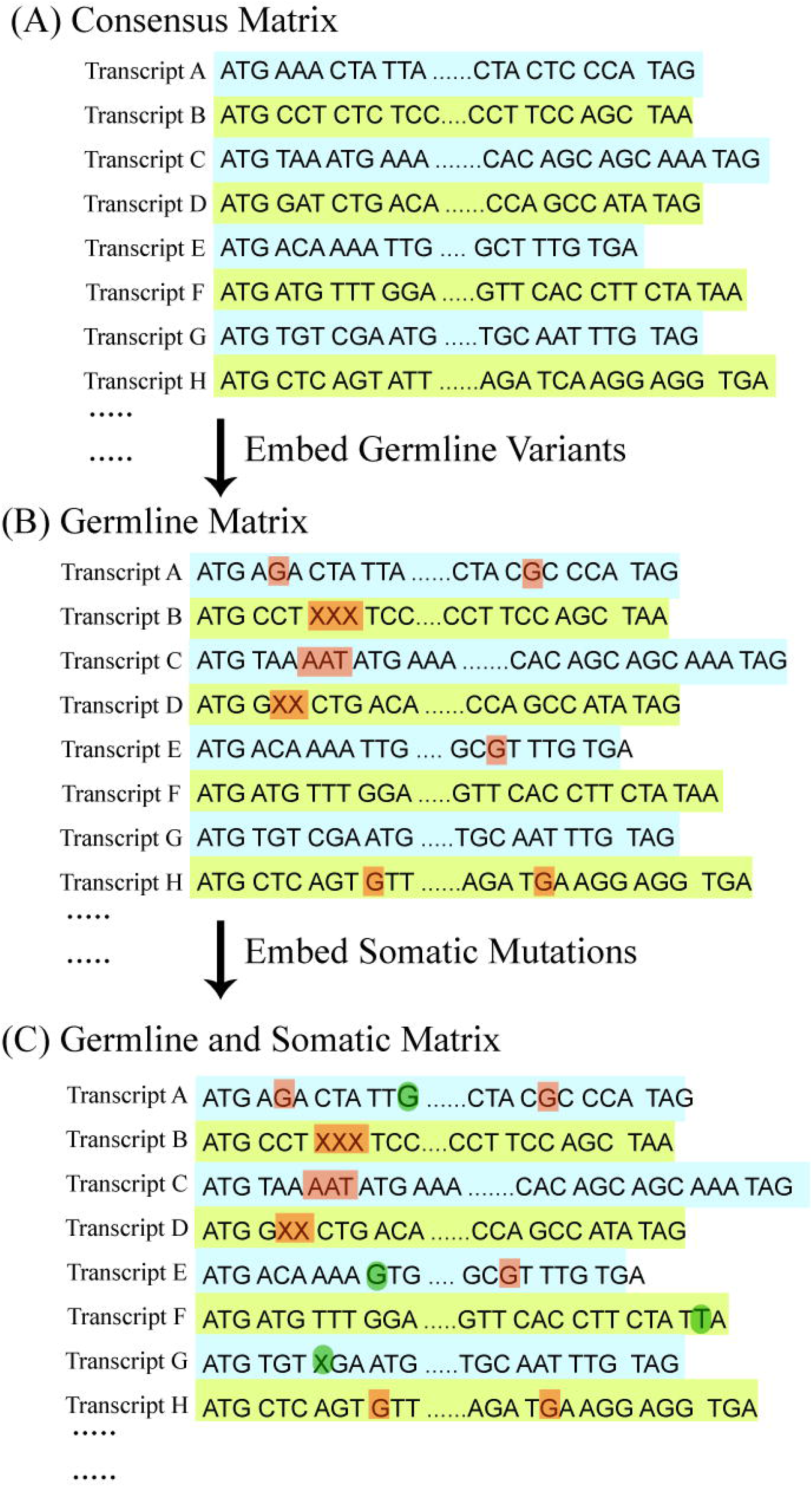
Feature generation for proposed models (A) The transcript sequences were retrieved from RefSeq and were formed as a consensus matrix (B) Each patient’s germline variants were embedded in the consensus matrix, forming a germline matrix for each sample. The brown dots are the germline variants including polymorphisms, deletions, and insertions. As an illustration, single nucleotide polymorphisms were identified and embedded in transcript A, E, and H. An in-frame shift deletion was embedded in transcript B and an in-frame shift insertion was embedded in transcript C. A frame shift deletion and frame shift insertion is embedded in transcripts D and E, respectively. Transcript F and G remained the same. (C) Each patient’s somatic mutations were embedded in the germline matrix (from B), forming a germline and somatic matrix. The green dots are the somatic mutations including SNVs, insertions, and deletions. As an illustration, the tissue gained somatic mutations in transcript A and E; gained a stop loss in transcript F; and gained a deletion that shifted the frame in transcript G.

It has been suggested that mutations prefer certain codons and the distance between amino acid changes have been described (33). Moreover, the position within the codon where the mutation occurs will determine if the expressed mutation is nonsynonymous or potentially synonymous. To incorporate codon information into our model features, one hot encoding was applied with every three nucleotides and was encoded as a binary unit. The combination of four nucleotides (A, C, T, and G) results in a vector with 64 dimensions to represent each codon combination.

### Model structure

A convolutional framework that consists of multiple layers was used in our study (Figure 2). The framework has three components: input layer (Figure 2A), encoder layer (Figure 2B) (multiple convolutional and dense layers), and fully connected layer (Figure 2C). Due to the nature of 64 codons in human genetics, the input layer in component (A) uses one hot encoding to represent each input sequence as a N*64 binary matrix, where N equals the number of codons. Therefore, the input can be considered as a 1-D sequence with 64 channels. Component (B) is an encoder layer to encode the input to a lower dimensional vector. The encoder component contains a sequence of convolutional layers with six output channels and a fully connected layer for each output channel. Therefore, a vector of six outputs is generated by the encoder for each input sequence. In theory, the output channel can be set as any positive integer. The more output channel, the more expressive capacity and more complexity of the model. To make a trade-off between the complexity and the expressive capacity, we set the output channel as six. In fact, if the precision of each channel is 0.01 (i.e., can store 100 numbers), 6 channels can express 100^6^ different samples. One convolutional layer is composed of one 1-D convolutional layer followed by a Leaky Rectified Linear Unit (LeakyReLU) (34) as the activation function and an average pooling layer. The number of convolution layers is determined by the transcript length N and the kernel size for average pooling layer. A Kernel size of six was used for the average pooling. Therefore, we will have log_6_N convolution layers for each transcript. Component (C) is a fully connected layer with k outputs for k diseases. The inputs of component (C) are the combinations of products from the component (B) generated under the sequence of transcripts. With the average of 3,375 bases in the transcripts, the encoder layer would have an average of 3∼4 convolution layers. To note, we set the following parameters for our model: the number of input channels for the encoder layer: 64; the convolution kernel size: 3, the output channel size of the encoder layer: 3; learning rate: 0.001; batch size: 32; number of learning epochs: 30. We used cross entropy loss as the loss function and Adam algorithm (35) as the optimizer.

**Figure 2:**
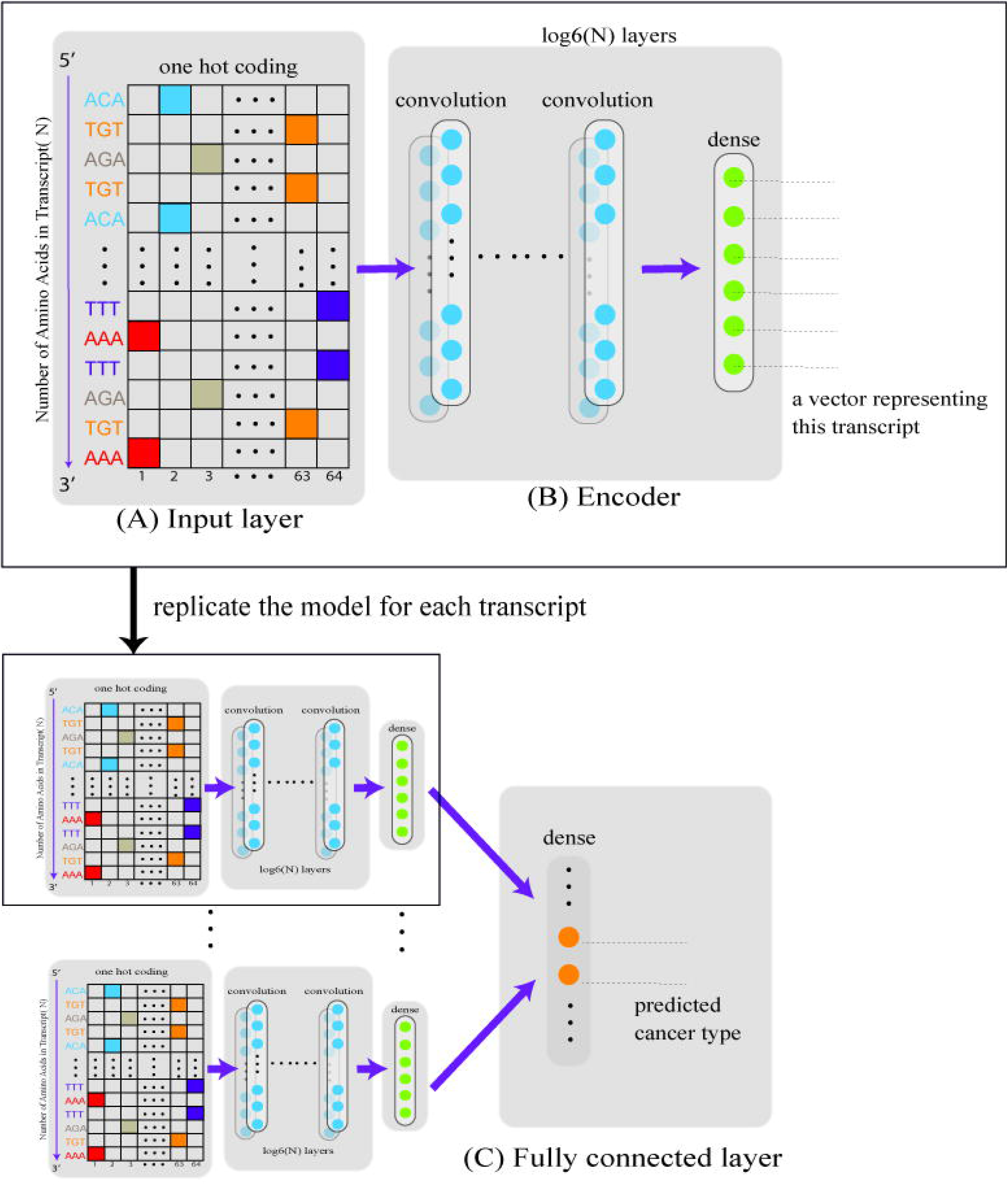
The architecture of the convolutional neural network. Component (A) is the input layer with one hot encoding with the column number equals 64 (number of total possible codons) and the row number equals the number of codons in the transcript. Component (B) is the encoder component containing a sequence of layers, each consisting of a convolutional layer, followed by a Leaky Rectified Linear Unit and average pooling layer. The number of convolution layers is determined by the gene length. Component (C) is a fully connected layer that combined all the outputs from the component (B) and has k outputs for k diseases

### Model evaluation and relevant gene discovery

A training set, validation set, and test set were created by randomly splitting the samples using a 7:1:2 ratio, respectively. Parameters were trained using the training set and tuned using the validation set. Precision, recall, and F-measure were calculated for each cancer type using the testing set. To compare the performance of our models to other conventional methods for cancer classification, we applied penalized logistic regression and linear support vector machine (SVM) (19,36). The performance was also compared between the germline matrix alone and the germline/somatic matrix. For the DeepCues, evaluations were repeated ten times with different initial seeds.

To reduce computational load, we selected genes that have been implicated in cancer using a list of 719 consensus genes (Table S1) from the Catalogue of Somatic Mutations in Cancer (COSMIC). COSMIC is a mutation catalogue with comprehensive mutation information curated from about 542,000 tumor samples (37). In our dataset, we found these consensus genes corresponded to 985 transcripts (Table S2) and used these transcripts to train and evaluate classifiers. The baseline model was also trained using germline variants and somatic mutations found in these selected transcripts. To discover potentially relevant genes not known to be implicated in cancer, we also applied a multinomial logistic regression model to the remaining transcripts using disease type as an output, and the number of mutations in each transcript as inputs to identify the 985 top ranked transcripts based on p-value (Table S3). Classifiers were trained, and evaluation was measured using only known pathogenic transcripts and also using a combination of the known and unknown pathogenic transcripts. It has been demonstrated that features frequently ranked high in different training sets yields a robust set of predictive features with stability (38). To obtain a gene list with reasonable stability, we repeated training the classifiers with random seeds and reported the top 20 most frequent transcripts in each replication.

## Results

### Classifier performance using known pathogenic transcripts

We first trained convolutional neural networks (CNNs) using the 985 known pathogenetic transcripts and calculated overall accuracy for all seven cancer types. Using only the germline matrix as an input, we achieved an overall accuracy of 73.9% (SE=0.7%) (SE is standard error). Using the germline/somatic matrix as an input, we achieved an overall accuracy of 77.6% (SE=0.9%). To compare our method with other conventional cancer classification methods, we calculated baseline accuracies using logistic penalized linear regression and linear SVM. Logistic penalized linear regression resulted in an overall accuracy of 51.5% (SE=0.5%) and 65.5% (SE=0.3%) using the germline matrix and germline/somatic matrix, respectively. Linear SVM resulted in an overall accuracy of 49.4% (SE=0.4%) and 58.6% (SE=0.3%) using the germline matrix and germline/somatic matrix, respectively (Figure 3). We found that our method significantly (p=4.5E-25) outperforms these methods.

**Figure 3:**
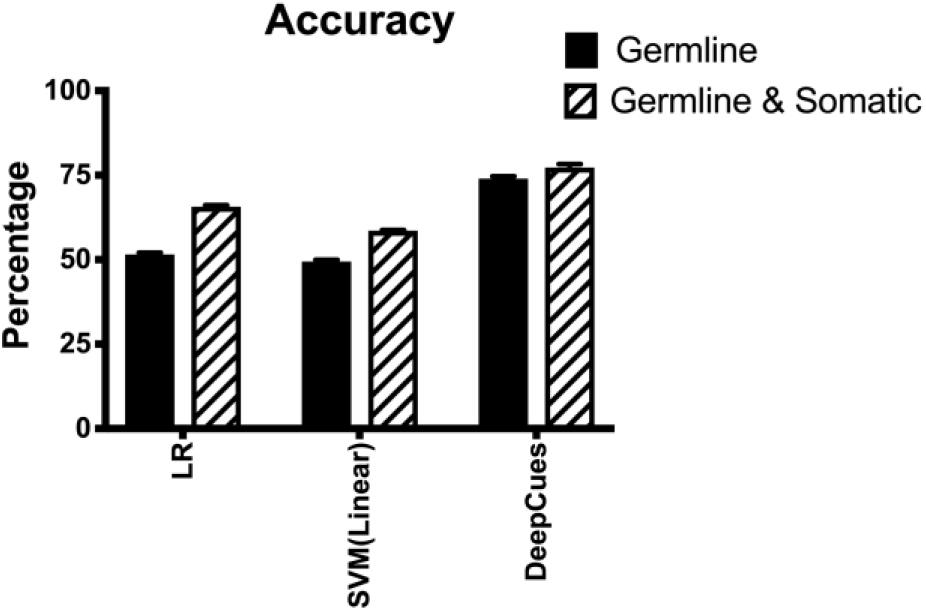
Comparing Prediction accuracy between DeepCues and baseline models, including penalized logistic regression (LR) and support vector machine (SVM) with linear kernel.

We performed classification for seven types of cancer: brain cancer, breast cancer, colorectal cancer, kidney cancer, lung cancer, prostate cancer, and uterus cancer. For each type of cancer, we calculated precision, recall, and f-measure using either the germline matrix or the germline/somatic matrix (Table 1). Using only germline data, we found breast cancer and colorectal cancer had the highest F-measure scores. Using both germline and somatic mutation data, we found breast cancer, colorectal cancer, and brain cancer had the highest F-measure scores. Adding the somatic mutation data, F-measures for breast cancer, brain cancer, and uterus cancer increased significantly (p=6.7E-03, 3.8E-06, and 1.9E-02 respectively).

### Classifier performance using known and unknown pathogenic transcripts

Similarly, to the prior analysis, we applied the same model and integrated both the 985 known and the 985 unknown pathogenic transcripts. Using only the germline matrix as an input, we achieved an overall accuracy of 82.68% (SE=0.6%). Using the germline/somatic matrix as an input, we achieved an overall accuracy of 80.0% (SE=0.9%).

We performed classification for the seven types of cancer. For each type of cancer, we calculated precision, recall, and F-measure using either the germline matrix or the germline/somatic matrix (Table 3). Using only germline data, we found breast cancer and colorectal cancer had the highest F-measure scores. Using both germline and somatic mutation data, we found breast cancer, colorectal, and uterus cancer had the highest F-measure scores.

**Table 2:**
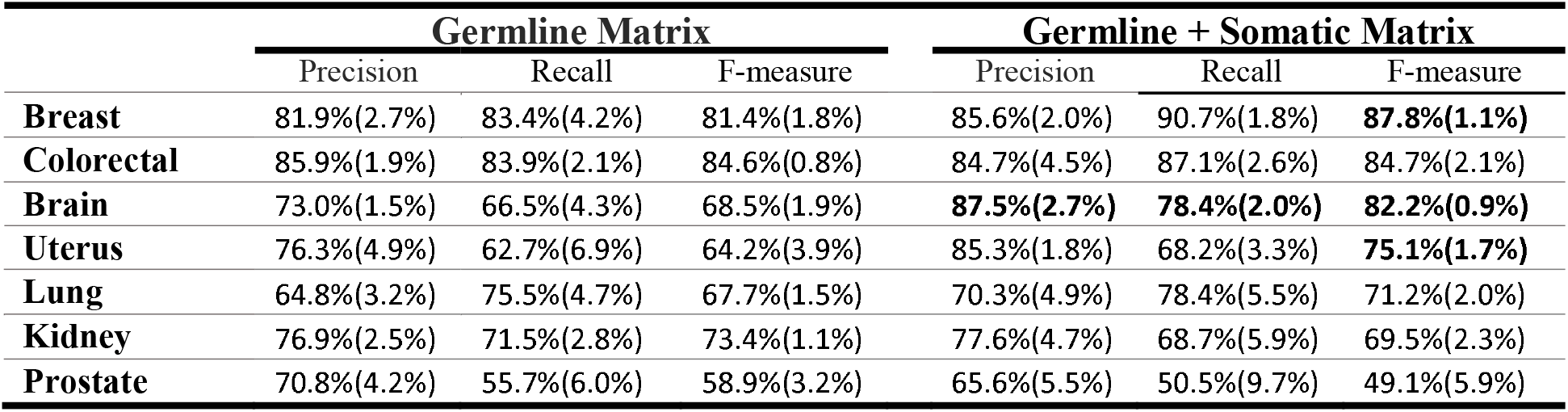
Precision and recall for our proposed model. The experiment is replicated for 10 times and the number in parenthesis is standard error. The bolded number are those that significantly improved after adding somatic mutation information.

**Table 3:**
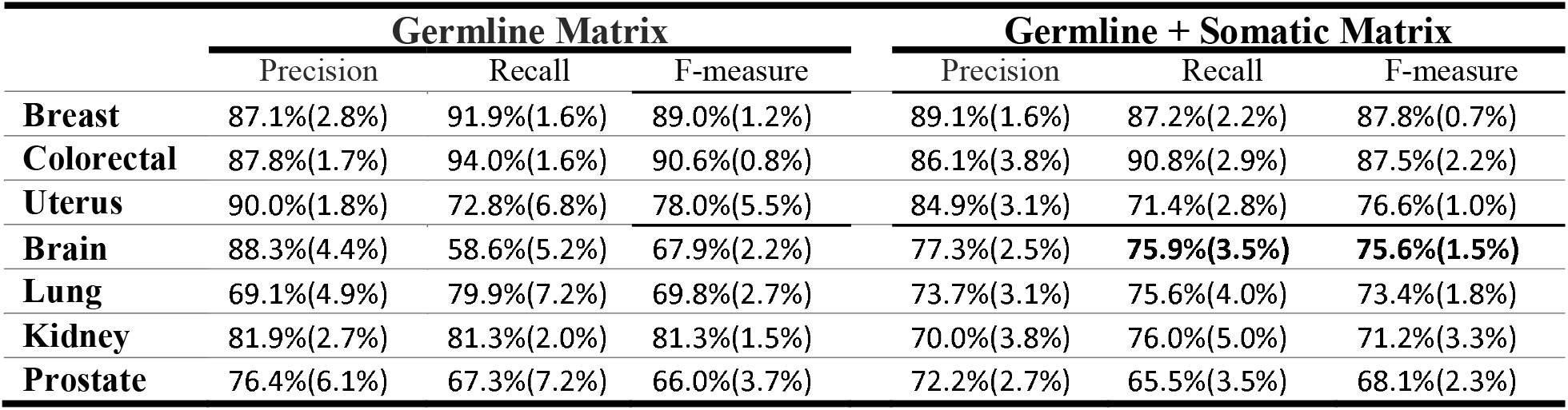
Precision and recall for our proposed model. The experiment is replicated for 10 times and the number in parenthesis standard error. The bolded number are those that significantly improved after adding somatic mutation information.

### Relevant genes discovery

Using the coefficients derived from the fully connected layer, the model can be extended to prioritize genes that are relevant for each cancer type. The analysis was repeated 10 times with different initial seeds and the top 20 genes were selected in each replicate. The genes were then ranked by appearance frequency in all the replicates. We performed relevant gene discovery for breast cancer and reported the top 10 for each study. We identified genes using the germline matrix alone and the germline/somatic matrix using the known pathogenic transcripts (Table 4) and also the known/unknown pathogenic transcripts (Table 3). We also performed relevant gene discovery for the other cancers using only the germline matrix (Table S3;S5) and the germline/somatic matrix (Table S4;S6). Using Germline/Somatic genes, 8 of the top 20 genes overlap with the COSMIC top 20 genes for breast cancer. The high consensus rate (40%) validated that our method is effective in identifying the relevant genes. We have also identified relevant genes that are unknown for breast cancer (bold genes under the panel of 1970 transcripts; Table 4).

**Table 4:**
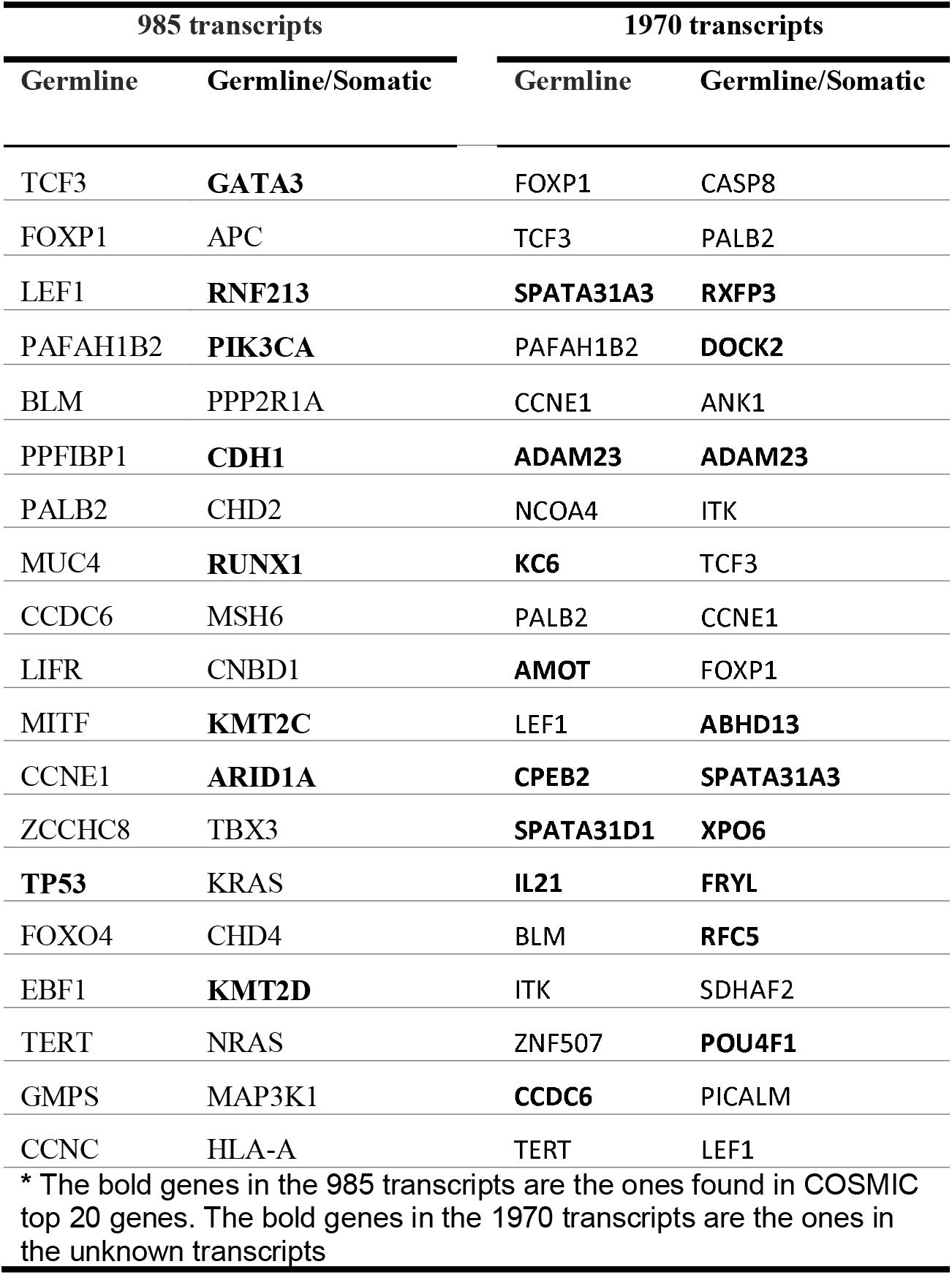
The top 20 genes relevant genes with breast cancer derived from the 985 pathogenetic transcripts and the 1970 transcripts.

## DISCUSSION

The development of high throughput sequencing technology has enabled the cataloging of large-scale genetic information. To help improve cancer diagnosis and targeted therapies, cancer type classification methods are continually being upgraded. Traditionally, the majority of classification methods based on DNA sequencing data has relied on studying single point somatic mutations with various regression models (15,39,40). Mutations involving insertions and deletions as well as germline mutations have been largely ignored due to the high dimensionality problem. Given that many methods are already limited in their ability to study so many variables, it has been even more challenging to integrate these variables and study them interactively. To deal with these challenges, groups have proposed aggregating mutations on a gene level to be studied as a feature (39,41,42). Mutations within genes have also been proposed to be studied within a matrix as inputs for machine learning methods (40,43). In our study, we have proposed a novel method, DeepCues. DeepCues integrated all somatic mutations and germline variants, including INDELs, to be studied as inputs in an joint manner. Convolutional Neural Networks (CNNs) were then applied to train classifiers for cancer type classification. Furthermore, we have included a fully connected layer to allow for relevant gene discovery to help characterize genes and pathways important for multiple cancers.

As a use case, we retrieved germline and somatic DNA sequencing data from matched samples across seven types of cancer and used DeepCues to perform cancer type classification. Using 985 known pathogenic transcripts as an input, we obtained 73.9% and 77.6% accuracy when using germline data alone and germline/somatic data, respectively. Following the integration of additional 985 unknown pathogenic transcripts into the model, we were able to increase overall accuracy to 82.7% and 80.0% for germline data and germline/somatic data, respectively. DeepCues was found to significantly outperform conventional methods used to perform cancer type classification (p=4.5E-25). Consistent with somatic mutations playing a large role in cancer (44), integration of somatic mutations together with germline data significantly improved overall accuracy (p=0.005) using the 719 known consensus genes. Integration of somatic data significantly increased accuracy for breast cancer (p=6.6E-03), brain cancer (p=3.8E-06), and uterine cancer (p=1.92E-02), suggesting somatic mutations play a relatively larger role in these cancers. Conventional methods have been limited regarding germline variation and their interactive role in cancer due to a large number of variables and complexity issues. In our study, we were able to obtain reasonable accuracy performances using the germline matrix only as an input. This suggests that germline variation may be more important than previously reported based on prior methods (15). More specifically, we found that breast cancer and colorectal cancer have the best performance using only germline information, suggesting that these two cancers probably confers higher heritability compared to others. Studies have reported high familial heritability in breast cancer and colorectal cancer too (45). Using a fully connected layer in our framework, we identified relevant both known and unknown pathogenic genes found using both the germline and germline/somatic data. For the 20 genes we have identified to be relevant for breast cancer, 40% of the genes have been reported in the COSMIC top 20 genes for breast cancer.

Future development to better evaluate and assess our model will involve the inclusion of gene expression level, copy number variation, methylation, as well as including additional transcripts to be studied. Given that DeepCues is novel in its ability to utilize germline data in an informative manner, it will be of great interest and clinical impact to apply DeepCues to differentiate cancerous and non-cancerous samples. Disease classification not only allows for improved diagnosis and therapies but also allows research to understand a disease through identified groups of genes and related pathways. DeepCues uses genetic sequencing data as inputs with little domain knowledge and feature preparation. With the abundance of genomic information available, we expect DeepCues can be used in a variety of disease settings to help profile diseases.

## SUPPLEMENTARY DATA

Table S1

Table S2

Table S3

Table S4

Table S5

Table S6

## Acknowledgement

We acknowledge the Center of Genomics Compute Cluster in Northwestern University for offering the computational resources and technical supports

